# How old are dragonflies and damselflies? Odonata (Insecta) transcriptomics resolve familial relationships

**DOI:** 10.1101/2020.07.07.191718

**Authors:** Manpreet Kohli, Harald Letsch, Carola Greve, Olivier Béthoux, Isabelle Deregnaucourt, Shanlin Liu, Xin Zhou, Alexander Donath, Christoph Mayer, Lars Podsiadlowski, Ryuichiro Machida, Oliver Niehuis, Jes Rust, Torsten Wappler, Xin Yu, Bernhard Misof, Jessica Ware

**Author notes:** corresponding author, **Corresponding authors contact information:** Dr. Manpreet Kohli, Dr. Harald Letsch.

## Abstract

Dragonflies and damselflies, representing the insect order Odonata, are among the earliest flying insects with living (extant) representatives. However, unravelling details of their long evolutionary history, such as egg laying (oviposition) strategies, is impeded by unresolved phylogenetic relationships, an issue particularly prevalent in damselfly families and fossil lineages. Here we present the first transcriptome-based phylogenetic reconstruction of Odonata, analyzing 2,980 protein-coding genes in 105 species representing nearly all of the order’s families (except Austropetaliidae and Neopetaliidae). All damselfly families and most dragonfly families are recovered as monophyletic groups. Our Molecular clock estimates suggest that crown-Zygoptera (damselflies) and -Anisoptera (dragonflies) both arose during the late Triassic. Several of the observed long inner branches in our topology are indicative of the extinction of once flourishing lineages. We also find that exophytic egg laying behaviour with a reduced ovipositor evolved in certain dragonflies during the late Jurassic / early Cretaceous. Lastly, we find that certain fossils have an unexpected deterring impact in divergence dating analysis.

## Results and discussion

Dragonflies and damselflies (Odonata) comprise three predatory suborders, Anisozygoptera, Zygoptera and Anisoptera, which are ubiquitous in lentic (flowing) and lotic (still-water) habitats [1]. Habitat choice in dragonflies are closely related to their oviposition behaviour. Not all dragonflies lay their eggs in the same fashion: there is the faster squirting-style exophytic oviposition, versus the slower endophytic oviposition where the eggs are laid inside plant material [2]. Egg laying can be risky as odonates are vulnerable while at the water and are routinely consumed by predators like fish, frogs and birds while mating and egg laying. Therefore, origin of these predators may have influenced oviposition behaviour in odonates. However, the evolution of egg-laying strategies is unclear due to a lack of resolution in the Odonata phylogenetic tree (e.g., [3, 4]). To understand the evolution of these odonate life history traits we used transcriptome sequences from 105 dragonfly and damselfly species along with a comprehensive fossil dataset (using newly assessed fossils in combination with those from Kohli *et al.* [5] (Supplementary Table S1, Table S7, Figure S3) to produce a well resolved time calibrated phylogeny of Odonata (Figure 1 and 2). The tree topologies we recovered using amino acids, and two nucleotide datasets (AA, Figure 1; NT2, NT123 Supplementary Figures S1 and S2) are congruent with respect to suborder monophyly and interfamilial relationships. All branches were recovered with maximal statistical non-parametric bootstrap support with the exception of two clades, within Anisoptera (Figure 1).

**Figure 1.**
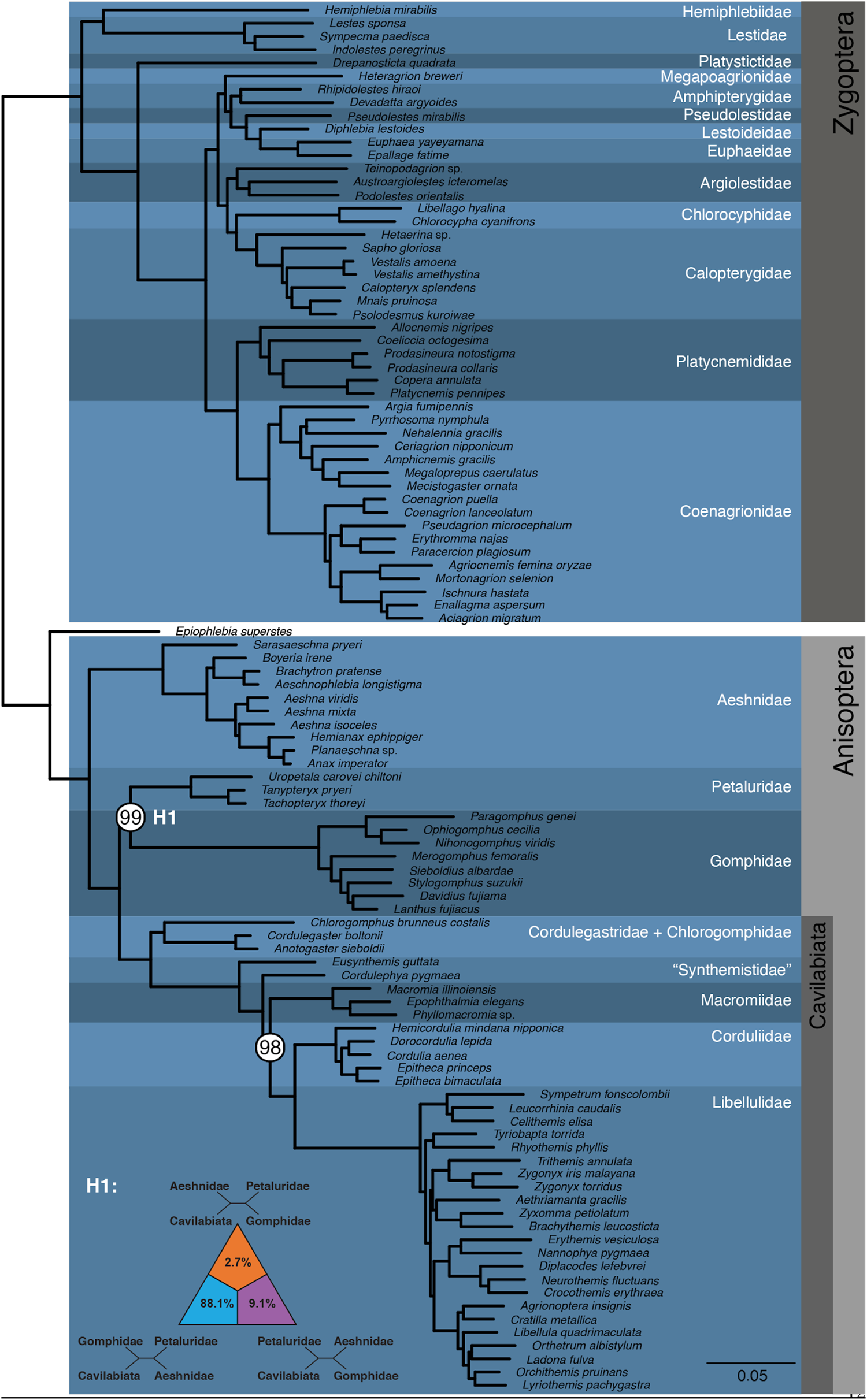
**Phylogenetic relationships in Odonata recovered** using the amino acid (AA) dataset. All the branches are recovered with 100% bootstrap support except the two nodes in Anisoptera highlighted with white circles on the nodes. Results of the four-cluster likelihood mapping (FcLM) for relationships among Petaluridae, Gomphidae and Cavilabiata are also represented.

**Figure 2:**
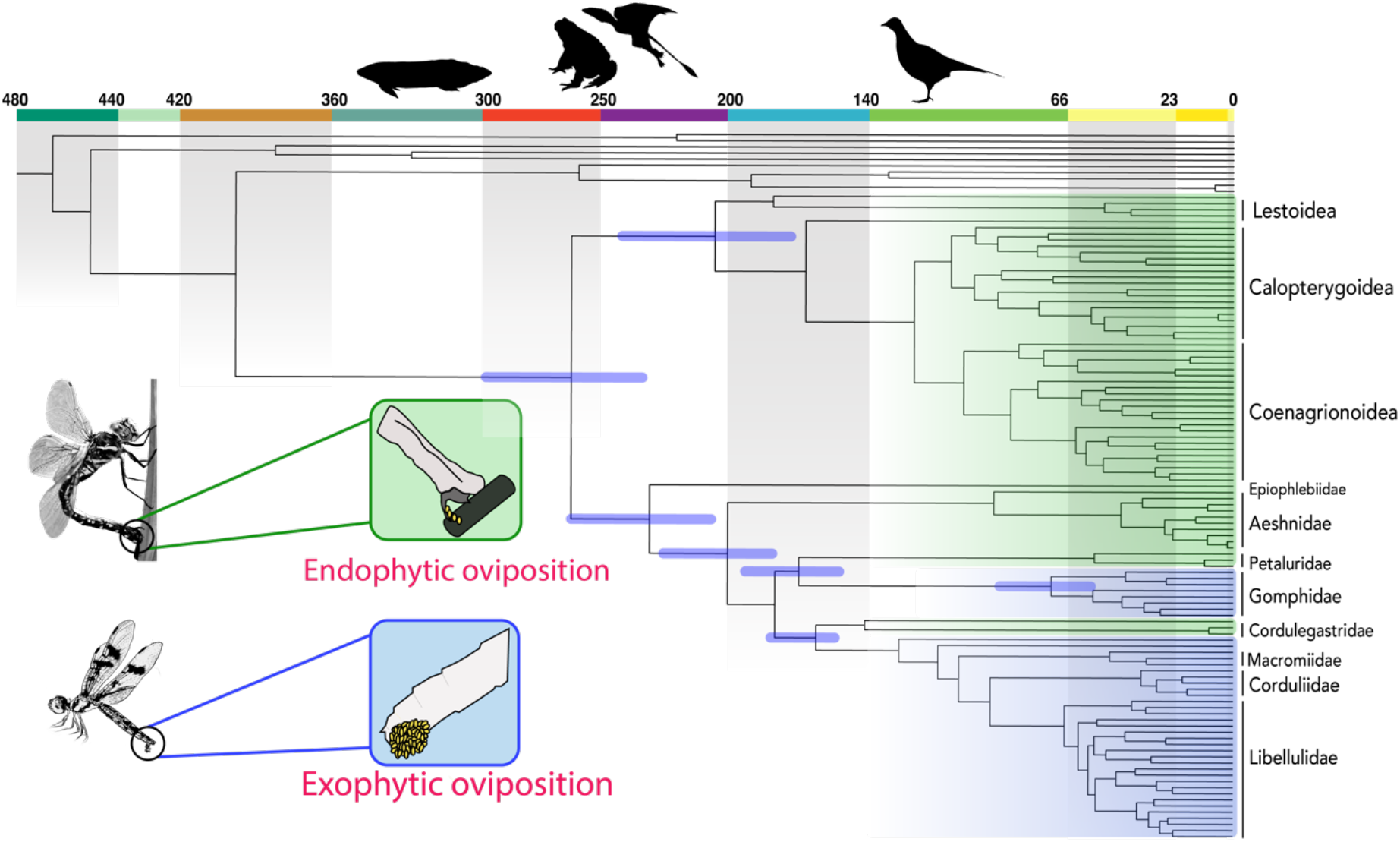
**Evolution of egg-laying behaviour in Odonata** are shown in accordance to the recovered phylogenetic relationships. Lineages with endophytic oviposition, those that lay their eggs in plant material, are shaded in green. Lineages with exophytic oviposition (usually laying eggs on the surface of water) are shaded in blue. Origin of the predators that may have influenced the evolution of egg-laying strategies in dragonflies and damselflies are represented along the geological time scale. Silhouettes of putative predators that were present at various geological times (left to right): fish, frogs, pterosaurs and birds. All the ages along the geological time scale are in millions of years.

Odonata are recovered as an ancient lineage, which probably originated during the Carboniferous. Indeed, the fossil record suggests that during the Pennsylvanian (323–298 Mya) and the Permian (298–251 Mya), stem-odonates were both diversified and ubiquitously distributed, including often famously large griffinflies, as well as the more gracile, damselfly-like †Archizygoptera [6-8]. Our results suggest that the crown-Odonata, dragonflies and damselflies as we recognize them today, diverged from their ancient relatives during the Permian independent of the fossil calibrations strategy (Figure 3).

**Figure 3:**
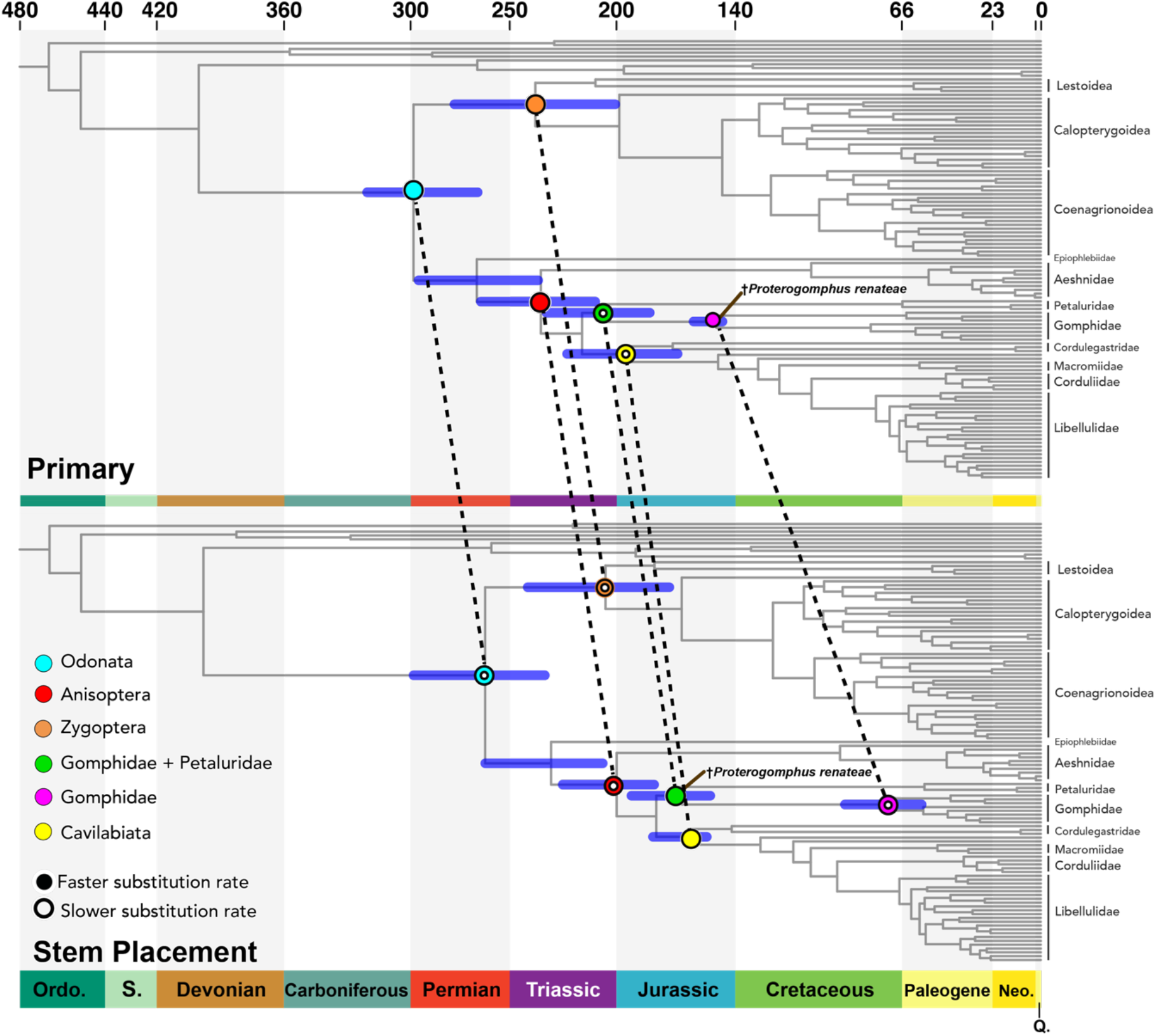
**Divergence times estimates comparisons** between crown or stem placement of the fossil *†Proterogomphus renateae* (indicated with brown line). Six different nodes of interest are indicated with different colors (cyan= Odonata, red=Anisoptera, orange=Zygoptera, green=Petaluroidea, pink=Gomphidae and yellow=Caviliabiata). These nodes are joined across the two scenarios with a dashed line to indicate the difference in the recovered divergence time estimates. We recover older ages under the *†Proterogomphus* crown placement scenario compared to the *†Proterogomphus* stem placement scenario. Hollowed circles on a node indicates a slower substitution rate compared to the same node in the other scenario. Substitution rates for the nodes Odonata, Anisoptera, Zygopter and Gomphidae were slower in the stem placement scenario compared to the crown placement scenario. By contrast, for nodes Zygoptera and Gomphidae + Petaluridae (Petaluroidea), rates in the stem scenario were faster compared to the crown scenario.

Damselflies (Zygoptera) originate around 206 million years ago (MYA) and are recovered as sister to Epiprocta, the group that comprises Anisozygoptera and Anisoptera (dragonflies) (Figure 1 and 2). In line with prior studies (e.g., [9-11]) *Hemiphlebia* is recovered as the earliest branching lineage of Lestoidea, which is in turn recovered as the earliest diverging lineage within Zygoptera. Superfamily Calopterygoidea, damselflies with often brightly coloured wings, consisting of families Chlorocyphidae, and Calopterygidae is recovered as 112 Myr old. Sister to Calopterygoidea, we recover the superfamily Coenagrionoidea, the bluets, which includes *Megaloprepus caerulatus* – odonate with the largest known wingspan [12]. Coenagrionidae are estimated to be 107 Myr old. Relationships in Zygoptera have previously been considered chaotic. Past studies have only resolved parts of the Zygoptera family tree [13-17] and a fully resolved phylogeny of Zygoptera didn’t exist until this present study.

Anisozygoptera, represented here by Japanese endemic *Epiophlebia superstes,* is recovered as an old group that originated in the mid Triassic (232 MYA) that is sister to the Anisoptera. Dragonflies as recognized today, i.e Crown-Anisoptera, emerged around 200 MYA. As has been shown in the past (e.g.,[1]), we indeed recover Aeshnidae as the earliest diverging lineage within Anisoptera. However, despite the vast amount of transcriptomic data used here, we still find a lack of resolution in Libelluloidea, and the interfamilial relationships among Gomphidae, Petaluridae and Cavilabiata (Figure 1). A group including Petaluridae and Gomphidae is recovered as sister to the superfamily Cavilabiata. Branch support for the sister group relationship of Petaluridae and Gomphidae is at 99%, a grouping that has been recovered previously, albeit with low support (e.g.,[11]). Yet, our four-cluster likelihood mapping (FcLM) does not provide support for this pair. FcLM analyses find Petaluridae+Gomphidae only supported by 11.7% of the quartets, whereas a clade comprising Gomphidae and Cavilabiata received high quartet support (78% of the quartets) (more details in the supplementary material). The grouping of Cavilabata and Gomphidae, has been found by other analysis [1], suggesting that the Cavilabiata and Gomphidae sister relationship may be difficult to resolve, perhaps due to short internodes, and/or rapid radiation within these groups. Further, a resolution is also confounded by the fact that Gomphidae seem to have much faster substitution rates than surrounding lineages nodes (suggested by long-branch leading to Gomphidae in Figure 1). Additional support for Gomphidae and Cavilabiata grouping is found in morphology. This group shares a reduction in the ovipositor for exophytic egg-laying. Ovipositor in damselflies, Aeshnoidea and Petaluridae comprises anterior and posterior gonapophyses enclosed by gonoplacs for endophytic oviposition in plant material (Figure 2). In Cavilabiata and Gomphidae, the gonoplacs are vestigial and in some Cavilabiata families the anterior and posterior gonapophyses are vestigial (see figures in [18]), suggesting the shared ancestor to Cavilabiata and Gomphidae possessed this reduction. The reduced ovipositor is used for exophytic oviposition, by spraying the eggs over the water surface. However, Matushkina [18] showed that some Cavilabiata retained ovipositor associated muscles and rudiments of the apparatus on the 9th abdominal segment, such muscles and rudiments were absent in the Gomphidae she examined. Since a reduction in the ovipositor seems to be different in the two exophytically ovipositing groups, this could suggest independent reductions. These independent reductions, however, don’t necessarily translate into support for a Petaluridae and Gomphidae sister relationship but rather they simply do not lend support for the Cavilabiata and Gomphidae sister relationship.

Gomphidae is estimated to have diverged in the Cretaceous (72 MYA), closer to the K/T boundary while Cavilabiata emerged during the late Jurassic (165 MYA). Based on these origin times Gomphidae and Cavalabiata are likely to have experienced different predation pressures. Several predators impose risks to odonates while undertaking oviposition: fish, frogs and birds commonly consume individuals that are copulating or ovipositing (Corbet, 1999). Frogs likely have been a strong source of predation pressure for a large portion of Odonate history as there are records of early Triassic frogs *(Triadobatrachus* [19]) [20]. Although the oldest crown birds emerged in the Cretaceous *(Asteriornis* [21]) there were likely stem birds before, hence it’s unclear how long they have been a source of predation on odonates. Bony fish have been around far longer, certainly predating the rise of Odonata, and presumably acting as a predation threat over the course of odonate evolution.

## A note on dating analysis

Fossil choice and placement have been shown to have an effect on recovered age estimates [13, 22-24]. Here, we particularly expected the inclusion/exclusion of the debated *↟Triassolestodes asiaticus* [25] to have the greatest effect on the node age estimates due to its placement on an ancient node, crown-Epiprocata, on the Odonata tree. On the contrary, we find that removal of this fossil neither changes the recovered ages, nor does it affect the precision of ages (Supplementary Figures S4–1,2 and Supplementary Table S7, under “Minus Triassolestidae” Scenario). The only observable differences, albeit small, are seen on Epiprocta (where the fossil calibration is placed) and its three surrounding nodes: Odonata, Zygoptera and Anisoptera. Out of these, crown-Odonata changed by 6.23 My (a 2.09% change), the largest change in node age seen throughout the tree. Epiprocta, the node from which the Triasssolestidae fossil was removed, itself only changed by 1.92%. Removal of *↟Triassolestodes asiaticus* led to increased precision, i.e, smaller confidence interval (CI) length on node age estimates, however, just like node ages, this change was very small on most nodes. The most noticeable impact of *↟Triassolestodes asiaticus* on the CI length was limited to the nodes Odonata (CI was reduced by 34.23%) and Epiprocta (CI reduced by 19.8%). Lastly, removal of *↟Triassolestodes asiaticus* didnot lead to dramatic changes in recovered substitution rates at nodes of interest (Supplementary Table S7). These results lead us to conclude that *↟Triassolestodes asiaticus* has a very localized effect. A more global impact of this fossil is perhaps prevented by the fact that all the nodes surrounding it also have fossil calibrations associated with them. However, in a scenario where the surrounding nodes are not calibrated, the impact of *↟Triassolestodes asiaticus* would perhaps be more profound. Our findings, however, do not imply that fossil choice is not pertinent to divergence time estimation. Rather, fossil choice cannot be treated separately from fossil placement as we discuss below.

We tested the impact of the placement of *†Proterogomphus renateae,* as this fossil was given a crown Gomphidae placement by Kohli *et al.* [5] but, we found it to be not an appropriate placement for the phylogeny recovered here due to taxon sampling. Choice of Gomphidae specimen used here for transcriptome sequence was in part based on specimen availability, and at the time of our sampling no Gomphidae molecular phylogeny had been reconstructed. Ware et al. [26] published a Gomphidae phylogeny after our sequencing was completed and revealed that the earliest branching lineages of Gomphidae were in the Lindeniinae, a lineage which we had not included here. Because of the absence Lindeniinae from our phylogenetic reconstruction *†Proterogomphus renateae* should not be placed on crown-Gomphidae as suggested by Kohli *et al.* [5]. Rather *†Proterogomphus renateae* is more appropriately placed on Gomphidae + Petaluridae node.

Unlike the impact of *†Triassolestodes asiaticus,* a stem placement of †*Proterogomphus renateae* led to strongly altered and younger ages on nodes throughout the topology (Figure 3). The largest age difference was seen on the crown-Gomphidae node which was recovered 81 million years younger (a 53% decrease in age) under the “Stem placement” scenario (Figure 3, stem placement scenario). Deeper nodes, such as Odonata, Epiprocta, Anisoptera, Zygoptera, and Cavilabiata were all recovered on average to be 30 million years younger (a 11–15% decrease in age) when *†Proterogomphus renateae* was used as a stem rather than a crown calibration. CI lengths either increased or decreased depending on the nodes. However, the absolute change in lengths of confidence intervals was high in several cases: on the Gomphidae node, the CI changed by 165%, increasing from 13.69 Myr to 36.35 Myr in breadth. An alarming aspect of the dramatic influence of this particular fossil is the extremely tight confidence interval recovered for crown-Gomphidae in the “Primary” scenario (Figure 3), which should be treated with extreme caution given the contrasting Stem placement results. In general, a short CI length has often been thought to imply a more accurate node age [27]. Hence, an extremely small confidence interval, like the one on crown Gomphidae, could mistakenly be interpreted as an extremely reliable result. However, the only reason we recover such a small CI for this node is because the age on that node is being pushed towards its upper limit with the older fossil calibration on it, while molecular data suggests a younger age on that node. Lastly, as with node ages and CIs, we see dramatic differences in substitution rates on branches leading up to certain nodes when *↟Proterogomphus renateae* is included or excluded (Supplementary Table S7). However, unlike node age and CI, the greatest impact of using *↟Proterogomphus renateae* as a stem calibration was seen on the branch leading up to Odonata, which made the substitution rate 24% lower compared to the primary scenario.

The impact of the gomphid calibration, underscores the importance of considering taxon sampling during the experimental setup phase, if divergence estimation of a group is an end goal. In fact, we found that several fossils which met the calibration vetting principles set forth by Parham *et al.* [28], were found to be unusable here simply because of our taxon sampling. Further, our iterative analyses using varying fossil calibration sets strongly highlight the relatively local rather than global impact of the fossil calibrations and underscores the importance of considering stem and crown placements of fossils when conducting divergence time estimation analysis. While some of the observed temporal gaps among clades may be due to our extant species taxon sample (i.e., perhaps due to undersampling the basal nodes of Gomphidae, for example), most ages are congruent with conclusions based on the fossil record. Moreover, several temporal gaps compose a genuine documentation of the extinction of once flourishing lineages. Crown-Odonata and crown-Aeshnidae, for example, should be regarded as only a small subset of their lineage’s historical diversity, which is reflected in our analysis as substantial temporal gaps. Other groups, such as Epiophlebiidae and Petaluridae, seem to have remained relatively species-poor throughout their history. It must be emphasized that upper boundaries of observed time gaps should not be mis-interpreted as evidence for extinction events---stem odonates have existed at the same time as crown odonates. A vivid example is the stem-odonatan ↟Protomyrmeleontoidea, which first originated in the Permian but are represented as a recently derived sample from the Early Cretaceous [29, 30], i.e. more than 100 My later to the origin of crown-Odonata.

The timing of appearance of the main extant lineages of Odonata do not clearly relate to global events which shaped present organismic life. Notably, the group does not seem to have experienced important radiation events nor severe replacement of its constituents. A possible exception is a putative mid-Cretaceous diversification of Zygoptera, suggested by both our analysis and the content of Myanmar amber [31], which would then be concomitant with the latest record of the †Protomyrmeleontoidea, which, judging from their wing morphology, must have occupied a flight mode niche similar as that of non-Calopterygidae Zygoptera. On a more general note, the ecological niche of Odonata, which can be depicted as that of generalist top predators inhabiting freshwater areas and capable of extensive dispersal, might have rendered the group relatively immune to historical global changes. One of the causes of extinction of particular lineages may have been a strong preference for oviposition host-plant which themselves may have declined, such as Sphenophytes, possibly used by griffinflies to host their eggs [32]. Also, given the critical role of flight performances for foraging, predator avoidance and reproductive success, their obsolescence might have become detrimental for some lineages, but this remains difficult to evaluate yet. Regardless of the assumed historical resilience of odonates, the fast, human-induced rarefaction and homogenization of habitats suitable to them [33] represents a threat that has no ancient counterpart.

## Acknowledgements

We thank Daniela Bartel for making the ortholog set, and Robert Waterhouse for analytics support. We thank Ondrej Hlinka for bioinformatics assistance, and Karen Meusemann for running phylogenetic pre and main analysis and editing the manuscript. Thank you to Alexander Blanke for leading the initial Transodonata 1KITE-working group. We thank all who provided specimens Alex Blanke, Akira Nishida, Chris Manchester, Daichi Kato, Guenther Theischinger, Jeanne Wilbrandt, John Trueman, Julian Glos, Kai Schütte, Kaoru Sekiya, Karen Meusemann, Manfred Niehuis, Martin Kubiak, Matthew Gimmel, Melissa Sanchez-Herrera, Miai Sakai, Mika Sugimura, Mitsutoshi Sugimura, Ola Fincke, Osamu Tanabe, Ralph S. Peters, Reiner Richter, Richard A. B. Leschen, Rudolf Meier, Hiroshi Yokota, Sae Nomura, Tanja Ziesmann, Thomas Buckley, Toshiyuki Teramoto, William Kuhn, Xiao-Li Tong, Xin Yu, Xin Zhou, Yoko Watanabe, Yuta Mashimo, Dominic Evangelista, and Erik Svensson. We are grateful to M. Hämäläinen for assistance with access to the relevant literature and the U.K. Geological Survey. ON thanks Netta Dorchin for help preparing field trips to Israel and acknowledges the Israeli Nature and National Parks Protection Authority for granting permission to collect samples.

## Author Contributions

### Manpreet Kohli

Collected fossil calibration data, ran divergence time analyses, wrote original draft of the manuscript, Figures and tables

### Harald Letsch

Collected fossil calibration data, assisted with divergence time analyses, wrote original draft of the manuscript, Figures and tables

### Carola Greve

Ran phylogenetic pre- and main analyses, review and edited the manuscript

### Olivier Béthoux

Coordinated and contributed to the collection of fossil calibration data, assisted with divergence time analyses, wrote original draft of the manuscript

### Isabelle Deregnaucourt

Collected fossil calibration data

### Shanlin Liu

Sequenced transcriptomes, conducted transcriptome assembly, performed data curation

### Xin Zhou

Collected specimens, oversaw transcriptome sequencing and assembly, performed data curation

### Alexander Donath

Performed transcriptome analyses, contamination checks, NCBI sequence submission, developed tools for analyses, review and edited the manuscript

### Christoph Mayer

Performed transcriptome analyses, data partitioning, review and edited the manuscript

### Lars Podsiadlowski

Performed transcriptome analyses, contamination checks, review and edited the manuscript

### Ryuichiro Machida

Collected specimens.

### Oliver Niehuis

Collected specimens.

### Jes Rust

Collected fossil calibration data

### Torsten Wappler

Collected fossil calibration data, review and edited the manuscript

### Xin Yu

Collected specimens.

### Bernhard Misof

Collected specimens, oversaw transcriptome sequencing and assembly, performed data curation

### Jessica Ware

Collected specimens, assisted with divergence time analyses, wrote original draft of the manuscript, Figures and tables

### Funding

JLW would like to acknowledge NSF grants #1564386 and 1453147. ON acknowledges the German Research Foundation (DFG) for supporting field trips within Germany and to Israel (NI 1387/1-1). Transcriptome sequencing and data curation under 1KITE were supported by funds from BGI-Shenzhen through XZ.

